# The burden of deleterious variants in a non-human primate biomedical model

**DOI:** 10.1101/784132

**Authors:** Vasily Ramensky, Anna J. Jasinska, Sandeep Deverasetty, Hannes Svardal, Ivette Zelaya, Matthew J. Jorgensen, Jay Ross Kaplan, J. Mark Cline, Anastasia Zharikova, Susan K. Service, Richard K. Wilson, Giovanni Coppola, Nelson B. Freimer, Wesley C. Warren

## Abstract

Genome sequencing studies of nonhuman primate (NHP) pedigree and population samples are discovering variants on a large and rapidly growing scale. These studies are increasing the utility of several NHP species as model systems for human disease. In particular, by identifying homozygous protein truncating variants (hPTVs) in genes hypothesized to play a role in causing human diseases, it may be possible to elucidate mechanisms for the phenotypic impact of such variants through investigations that are infeasible in humans. The Caribbean vervet (*Chlorocebus aethiops sabaeus*) is uniquely valuable for this purpose, as the dramatic expansion of its population following severe bottlenecks has enabled PTVs that passed through the bottleneck to attain a relatively high frequency. Using whole genome sequence (WGS) data from 719 monkeys of the Vervet Research Colony (VRC) extended pedigree, we found 2,802 protein-truncating alleles in 1,747 protein-coding genes present in homozygous state in at least one monkey. Polymorphic sites for 923 SNV hPTVs were also observed in natural Caribbean populations from which the VRC descends. The vervet genome browser (VGB) includes information on these PTVs, together with a catalog of phenotypes and biological samples available for monkeys who carry them. We describe initial explorations of the possible impact of vervet PTVs on early infant mortality.

## INTRODUCTION

The difficulty in determining whether specific variants are deleterious, and in identifying the phenotypic effect of such variants, are critical limitations to the utility of whole exome or whole genome sequencing within clinical medicine. Here we describe a strategy to aid this determination, through examination of protein truncating variants (PTVs) that were initially identified in humans but that occur naturally in populations of the Caribbean vervet nonhuman primate (NHP) model. Of the putatively deleterious variant types, PTVs have been the most extensively investigated, because their potential functional impact is most obvious and because their association with specific phenotypes can help to reveal casual pathways of disease. In humans, much of what is known about the biology of PTVs derives from studies of populations whose demographic histories increase the likelihood of identifying homozygous carriers of such variants. Examples include population isolates with extensive consanguinities, such as the Old Order Amish (Strauss and Puffenberger 2009) or that have experienced recent rapid expansion from extreme bottlenecks, such as Finland (Lim et al. 2014; Locke et al. 2019). The advent of sequencing studies of large human cohorts has generated a robust pipeline of PTV discovery, which has led to the identification of a number of large-effect associations to diseases or disease-related traits (Dewey et al. 2016). However, most of the PTVs identified through such studies to date are present in the genomes of apparently healthy individuals (MacArthur et al. 2012), suggesting that their phenotypic impact, if any, may not be obvious (Rivas et al. 2015; Jagannathan and Bradley 2016).

Investigations in model systems have traditionally provided an avenue to help distinguish between damaging and benign PTVs in the ever-expanding catalogs of publicly accessible sequence variant databases (e.g. ExAC (Lek et al. 2016), Database of Essential Genes (Luo et al. 2014), ClinVar (Landrum et al. 2016) and OMIM (https://omim.org). However the genetic, morphological, and physiological distance from humans of the most widely used model systems has often limited the utility of this strategy. Several NHP research colonies are comprised of old world monkeys (OWM) that are the most closely related to humans of all model systems, are managed in well-established pedigrees and are characterized by relatively limited inter-individual variability in environmental exposures such as diet. Such colonies offer an opportunity to study PTV effects in homozygous or heterozygous states with a degree of control over experimental variables that far exceeds that of human studies that have examined PTV effects (Lim et al. 2014; Sulem et al. 2015; Narasimhan et al. 2016a, 2016b).

To our knowledge no studies have attempted to associate human PTVs, broadly, to observed NHP phenotypes, and there is little information, to date, on the extent of overlap between human and NHP PTVs. Among captive rhesus macaques, two studies have shown various degree of shared PTVs with human, depending on the types of variant filters deployed (Xue et al. 2016; Bimber et al. 2017). In six great ape populations, a range of PTVs was reported, with a strict focus on the stop gain classification (de Valles-Ibáñez et al. 2016). Sundaram et al. developed a deep neural learning algorithm to identify pathogenic variants in patients affected with rare diseases of unknown etiology, based on the prevalence of the variants in NHPs (Sundaram et al. 2018). By training this neural network on six NHP species (including chimpanzee), they generated a database of 70 million missense variants that they proposed warranted further interpretation and validation. While great apes are evolutionarily closer to humans than OWM, experimental (or even observational) studies of the phenotypic impact of PTVs are not generally possible.

We previously generated a robust collection of SNVs for the vervet, a species used as a model for several human diseases (Huang et al. 2015) and a tissue gene expression and eQTL catalog, thus far not available from other NHPs (Jasinska et al. 2017). We describe here a new vervet genetic resource, a genome browser that catalogs genome-wide PTVs from 719 monkeys from the Vervet Research Colony (VRC) extended pedigree. For each of these monkeys, the browser provides phenotypic data and an inventory of biological samples that are available to the community.

## RESULTS

### Protein-coding sequence variation

We have previously described the design of the studies that generated WGS data from both the VRC pedigree and from vervet populations in Africa and the Caribbean, including sampling and sequencing (Huang et al. 2015; Svardal et al. 2017). To recover variation in the extended vervet exome, defined as gene transcript exons expanded by +/-50 bp and padded by 1000-bp gene flanks, we undertook a two-step process of variant calling, phasing, and imputation in the 719 VRC WGS monkeys. This process uncovered 1,051,886 single nucleotide variants (SNVs) and 241,648 short indels (Table 1). Although most of the discovered variant sites are biallelic, 6% of sites harbor more than one alternative allele.

**Table 1.** Vervet variant annotation and variant counts across 719 sequenced animals. Acceptor or donor variants that are predicted disrupt transcript processing. Missense variants confer amino acid changes with unknown impact in protein-coding genes. Stop or gain variants disrupt the processing of full-length transcripts. Synonymous variants are base changes that don’t confer amino acid changes and are considered neutral. Single nucleotide varaints (SNVs) in coding exons; Upstream-1000, downstream-1000: variants in 1kbp gene flanking regions; 3-UTR, 5-UTR, intron: variants in gene untranslated regions and introns. Non-coding, SNVs and indels in non-coding genes.

| *Type* | *SNV* | *Indel* | *Total* |
| --- | --- | --- | --- |
| Acceptor | 520 | 604 | 1,124 |
| Coding exon indel | N/A | 9,930 | 9,930 |
| Donor | 800 | 507 | 1,307 |
| Downstream-1000 | 212,581 | 56,487 | 269,068 |
| Intron | 130,781 | 34,320 | 165,101 |
| Missense | 70,212 | N/A | 70,212 |
| Non-coding | 113,244 | 22,665 | 135,909 |
| Stop-gain | 1,416 | N/A | 1,416 |
| Stop-loss | 131 | N/A | 131 |
| Synonymous | 76,469 | N/A | 76,469 |
| Upstream-1000 | 242,610 | 65,035 | 307,645 |
| 3-UTR | 122,669 | 35,804 | 158,473 |
| 5-UTR | 80,453 | 16,296 | 96,749 |
| Total | 1,051,886 | 241,648 | 1,293,534 |

Variants predicted to truncate protein-coding genes (stop gain, or splice site donor and acceptor SNVs, frameshifting indels in coding exons all collectively termed PTVs; see Methods) have, on average, lower alternative allele count (AAC) values (median values ranging from 2-90) than other classes of variants (median values ranging from 176-226; Figure 1). The average number of autosomal PTVs per monkey varies depending on the filters applied. Considering only high quality called genotypes (GQ>=20), each monkey has on average 166 splice site or stop gain PTVs across all genes with known human orthologs (Table 2). These averages slightly exceed the 130 per monkey observed for comparable PTVs among 21 sequenced rhesus macaques of mixed ancestry (Indian, Chinese and hybrids) (Bimber et al. 2017).

**Table 2.** Average number of autosomal polymorphic PTV variants per monkey. Homozygous variants are given in parentheses. High quality genotypes: called (not imputed) genotypes with GQ>=20 as reported by GATK. Genes with human orthologs but no uncharacterized genes were counted here.

| *Classification* | *Stop-gain* | *Splice site* | *Coding exon indels* | *Total* |
| --- | --- | --- | --- | --- |
| All genes/genotypes | 312 (67) | 476 (111) | 921 (205) | 1,709 (383) |
| All genotypes/genes with human orthologs | 189 (40) | 238 (52) | 608 (144) | 1,035 (236) |
| High quality genotypes/genes with human orthologs | 82 (9) | 84 (10) | 216 (19) | 382 (38) |

**Figure 1.**
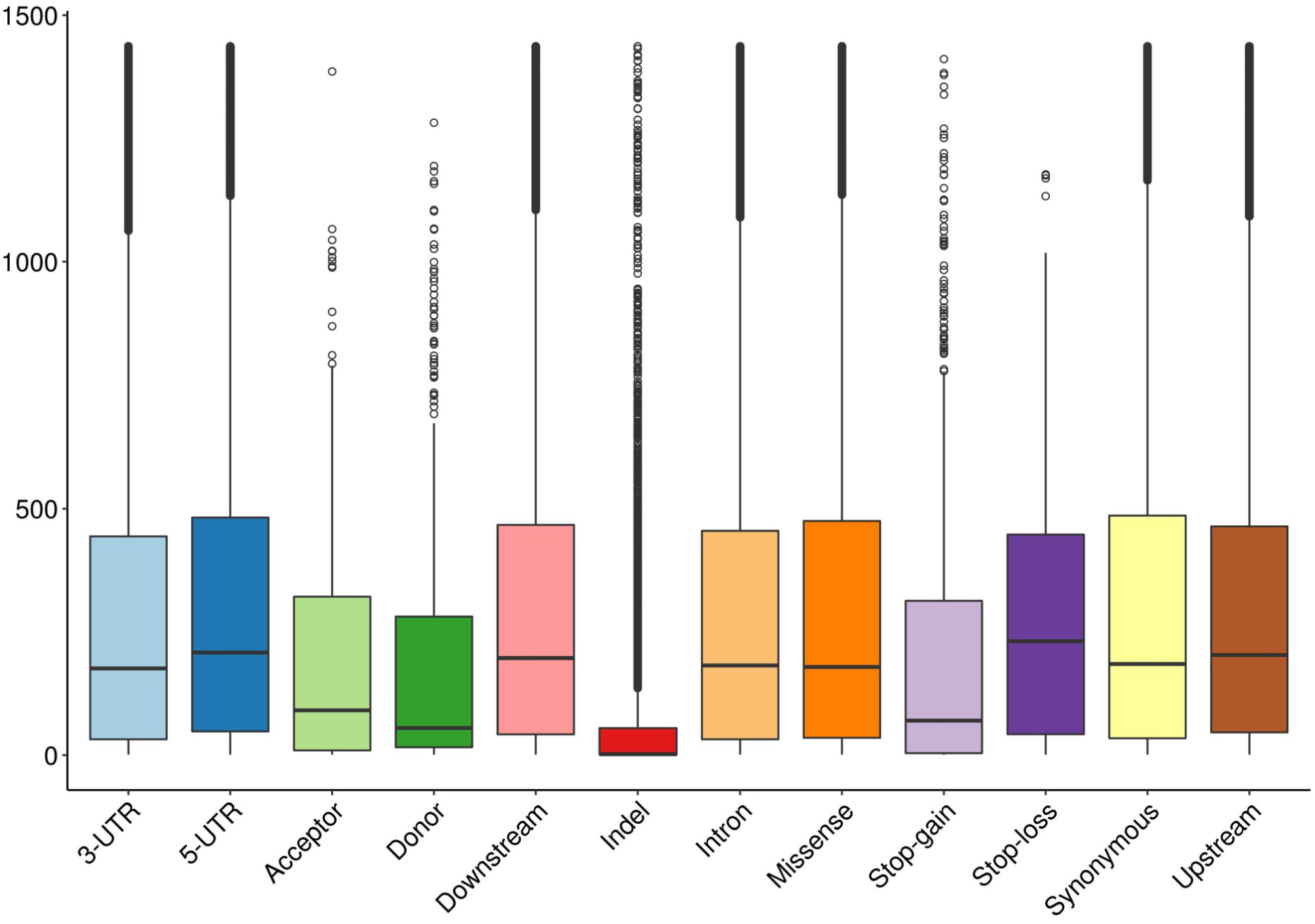
Alternative allele counts by variant type.

Polymorphic sites for 923 SNV homozygous protein truncating variants (hPTVs) were also observed in the natural Caribbean populations from which the VRC descends (Svardal et al. 2017). Supplementary Table 1 reports PTV alternative allele frequencies and numbers of monkeys with genotype calls in population samples.

### Vervet genome browser

The vervet monkey genome browser (VGB) is available at https://coppolalab.ucla.edu/vgb/home. The browser is built using JBrowse (https://jbrowse.org/) which is a fast genome annotation viewer (Buels et al. 2016). This browser allows a user to view objects such as genes and variants in their genomic context. The major view presents several tracks including NCBI RefSeq, ENSEMBL 1.82 and 1.91 gene annotation, PTV information, median gene expression RPKM values in 7 vervet tissues, and ChIP-Seq information (Supplementary Figure 1). A user starts by searching for a gene against the NCBI gene annotation track, and then proceeds to the major view which allows one to search against all available gene annotation tracks.

Using the genome browser, one can check what data and biological samples are available for monkeys harboring a specific PTV. Selecting a PTV from PTV-Short track displays a pop-up window with additional feature attributes such as position, gene, individual genotypes and genotype counts which are indicated as N_Het and N_HomAlt, links to phenotypic data, and an inventory of biological samples available for that PTV. Detailed phenotypic and biological sample availability for VRC monkeys, including a list of the 133 sequenced animals currently living in the colony, can be found in Supplementary Table 2.

### Segregating PTVs

The segregating vervet PTVs that we discovered, especially those present in a homozygous state, represent natural candidates for analyses aimed at identifying the phenotypic impact of vervet genetic variants. Out of 9,574 discovered PTVs in protein-coding genes, 2,802 alleles in 1,747 genes are present in a homozygous state in at least one vervet. Approximately 73% (2,047) occur in homozygous form in ≤ 10% of all sequenced vervets, however the remaining 27% are relatively common and exhibit a broad range of frequencies in the VRC population. There are 1,502 PTVs with at least one homozygous alternative carrier in 1,133 protein-coding genes with a known human ortholog (Supplementary Table 1). Also, compound heterozygotes that occur when single alleles of two different PTVs reside on different haplotypes of the same gene are found in 1,837 vervet genes harboring more than one PTV. Supplementary Table 3 contains individual genotypes and quality information for 2,802 homozygous PTVs in 133 currently living sequenced monkeys.

Among rare PTVs, we observed numerous genes implicated in risk for human diseases: Alzheimer disease (*PAXIP1, APBB2*), cancer (*CTNNB1, DLEC1, MSH2, TP53*), hypertension (*ECE1, NOS2*), Parkinson disease (*DNAJC13, VPS35, ATXN2, LRRK2*), mental retardation (*DEAF1, HERC2, SETBP1, CTNNB1, RPS6KA3, SHROOM4, SPATA5, ZDHHC15*), infectious diseases (*CD209, TIRAP*), Charcot-MarieTooth disease (*NDRG1*), Premature ovarian failure 8 (*STAG3*) (Köhler et al. 2014; Landrum et al. 2016)

### Expression of genes with homozygous PTVs

An earlier study reported a catalog of eQTL in vervets based on gene expression analysis in seven tissues of 58 sequenced VRC monkeys (Jasinska et al. 2017). To evaluate whether the vervet PTVs, overall, affect gene expression, we compared expression levels between monkeys with and without homozygous PTVs (hPTV), at all genes for which we observed at least one hPTV (Supplementary Table 4). For each such PTV we evaluated expression in each tissue separately and calculated regression coefficients (Betas) with accompanying significance levels for each hPTV-tissue combination that passed data quality filters. Negative Beta values indicated decreased expression in hPTV allele carriers. The initial test set included 8,683 PTV-tissue combinations for 1,518 hPTVs in 1,236 genes (Table 3). The stringent subset (see Methods) contains 3,918 combinations, of which 683 have nominal significance level P<0.05 with 393 showing decreased and 290 increased expression. If hPTVs are further split by types, the largest difference is observed for the splice site variants: 178 (66.7%) vs 89 (33.3%). We hypothesize that the observed association of hPTVs with a statistically significant increase in gene expression can be explained by their linkage with corresponding eQTL alleles. Homozygous PTVs negatively associated with expression may be linked with decreasing eQTL alleles or trigger nonsense-mediated decay with subsequent degradation of RNA directly. This may explain the observed prevalence of expression-decreasing hPTVs and agrees with the earlier observation that most predicted NMD-triggering variants have no detectable effect on gene expression (MacArthur et al. 2012).

**Table 3.** Association of homozygous protein-truncating variants with gene expression: negative or positive. The initial test set contained 8,683 PTV-tissue combinations. The stringent set (3,918) included only PTVs with at least three homozygous alternative and three homozygous reference genotypes. Negative (positive): decreased (increased) expression in homozygous PTV allele carriers. P-value reflects the probability that a PTV is not associated with expression, with p<0.05 cases considered as significant. Numbers in parentheses are % of total cases with p<0.05. The three bottow rows report the stringent set split by variant types.

|  | *Negative*<br>*$p < 0.05$* | *Positive*<br>*$p < 0.05$* | *Negative*<br>*$p \geq 0.05$* | *Positive*<br>*$p \geq 0.05$* | *Total*<br>*$p < 0.05$* | *Total* |
| --- | --- | --- | --- | --- | --- | --- |
| Initial | 623 (55.5) | 499 (44.5) | 3,898 | 3,663 | 1,122 | 8,683 |
| Stringent | 393 (57.5) | 290 (42.5) | 1,709 | 1,526 | 683 | 3,918 |
| Stringent, splice-site | 178 (66.7) | 89 (33.3) | 666 | 519 | 267 | 1,452 |
| Stringent, stop-gain | 63 (53.4) | 55 (46.6) | 346 | 331 | 118 | 795 |
| Stringent, coding indels | 152 (51.0) | 146 (49.0) | 697 | 676 | 298 | 1,671 |

### Homozygous PTV to phenotype association

As with all NHPs, there is currently no systematic catalog of extreme phenotypes in which to evaluate the impact of PTVs in vervet monkeys. Within the VRC early infant mortality is recorded. We hypothesized that hPTVs might play a role in this outcome (Lord et al. 2019) and found 29 homozygous and 5 compound heterozygotes that were present only in the 24 female monkeys with high rates of early infant mortality (high EIM monkeys, see Methods; Table 4, Supplementary Table 5). All such variants were present in a single monkey, except for a homozygous *Trp159Ter* stop-gain mutation in the *ARPC3* gene seen in two high EIM monkeys. After exclusion of 8 genes without known human orthologs (*LOCnnnnn* identifiers) and two olfactory receptors (*OR10A6, OR52Z1*), we investigated phenotype-related associations for the remaining 20 genes using gene-phenotype annotations from the Mouse Genome Informatics, Human Phenotype Ontology, OMIM, and ClinVar databases (see Methods). Nine of the 20 genes (*SLC25A36, SLFN13, THNSL2, TMPRSS9, IGHV2-5, TULP2, FAM65C, MTX1, NPEPL1*) have no annotation, whereas the remaining 11 genes (*ARPC3, RIPK3, SALL3, HBD, DNAJC16, MARCKSL1, MUS81, RAMP2, POLR2A, SULF1, ATXN2*) are annotated with an infant mortality relationship reported for each gene (Supplementary Tables 6 and 7). Two genes without annotations are said to be involved in protein-protein interactions with high EIM-related genes: *THNSL2* with *ARPC3* (MINT, IntAct) and *POLR2A* (STRING), *SLC25A36* with *SLC13A5* (STRING).

**Table 4.**
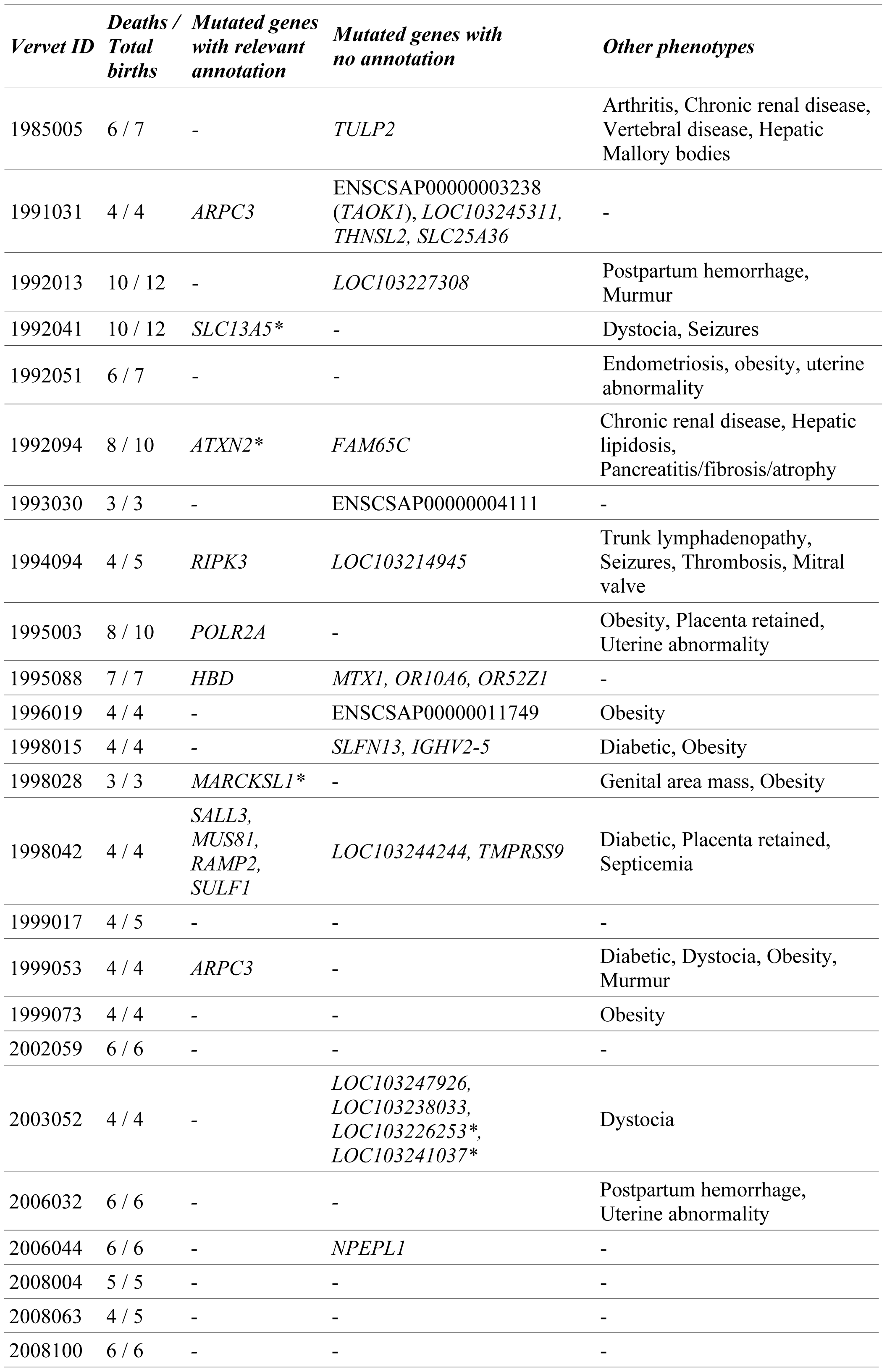
Genes with homozygous protein-truncating and compound heterozygous variants unique to 24 high early infant mortality vervets. Mutated genes with relevant gene ontology were aligned to observed phenotypes in the VRC. Genes with asterisk contain compound heterozygotes. The number of infants that were born live and died during the first month of life or were never seen alive due to stillbirths or miscarriage per female vervet is presented. If gene symbol is not known, protein id is provided (e.g., ENSCSAP00000003238).

We also present similar analyses for vervet hPTVs associated in humans with several other biomedically relevant traits including: cataract, electrocardiogram abnormality, hepatic Mallory bodies, malformations, and obesity with mass lumps (Supplementary Table 8). Among human or model organisms some candidate genes have been implicated in the same or related phenotypes. This is the case, for example, with respect to *WDR17* in cataract disease (Stöhr et al. 2002; Geisert et al. 2009) or *BID* in the human “cat eye syndrome”, characterized by congenital malformations and deformations (Footz et al. 2001), and in similar mouse organ morphology phenotypes (MGI:2158671). In other examples, a paralog of a vervet gene has been involved in a related phenotype, such as, *TULP1* which is associated with hepatic Mallory bodies or *SERPIN* family members which are implicated in cardio-related and multiple organ and bone morphology abnormality phenotypes (Supplementary Table 8).

### Human disease mutations in vervets

We also analyzed human pathogenic missense and stop-gain mutations from ClinVar that map to polymorphic sites in orthologous vervet genes (see Methods) and found 6 missense mutations for which phenotypes observed in vervets harboring alternative allele(s) can be related to clinical features reported in human patients (Table 5). In three cases (*PRSS1* p.Ala16Val, hereditary pancreatitis; *TRIM32* p.Pro130Ser Bardet-Biedl syndrome; *ACTB* p.Glu117Lys Baraitser-Winter syndrome) the human pathogenic allele matches the vervet alternative allele. Although the matched human alleles in three separate vervet genes are known to be highly penetrant in human, we only see related phenotypes in some vervets. The lack of manifestation in other vervets with a predicted penetrant disease allele may be due to the lack of systematic phenotyping of VRC monkeys, or because some of them did not reach the age when the symptoms are observed. It is also possible that this observation reflects variable penetrance resulting from the modulatory effect of cis-regulation (Castel et al. 2018).

**Table 5.**
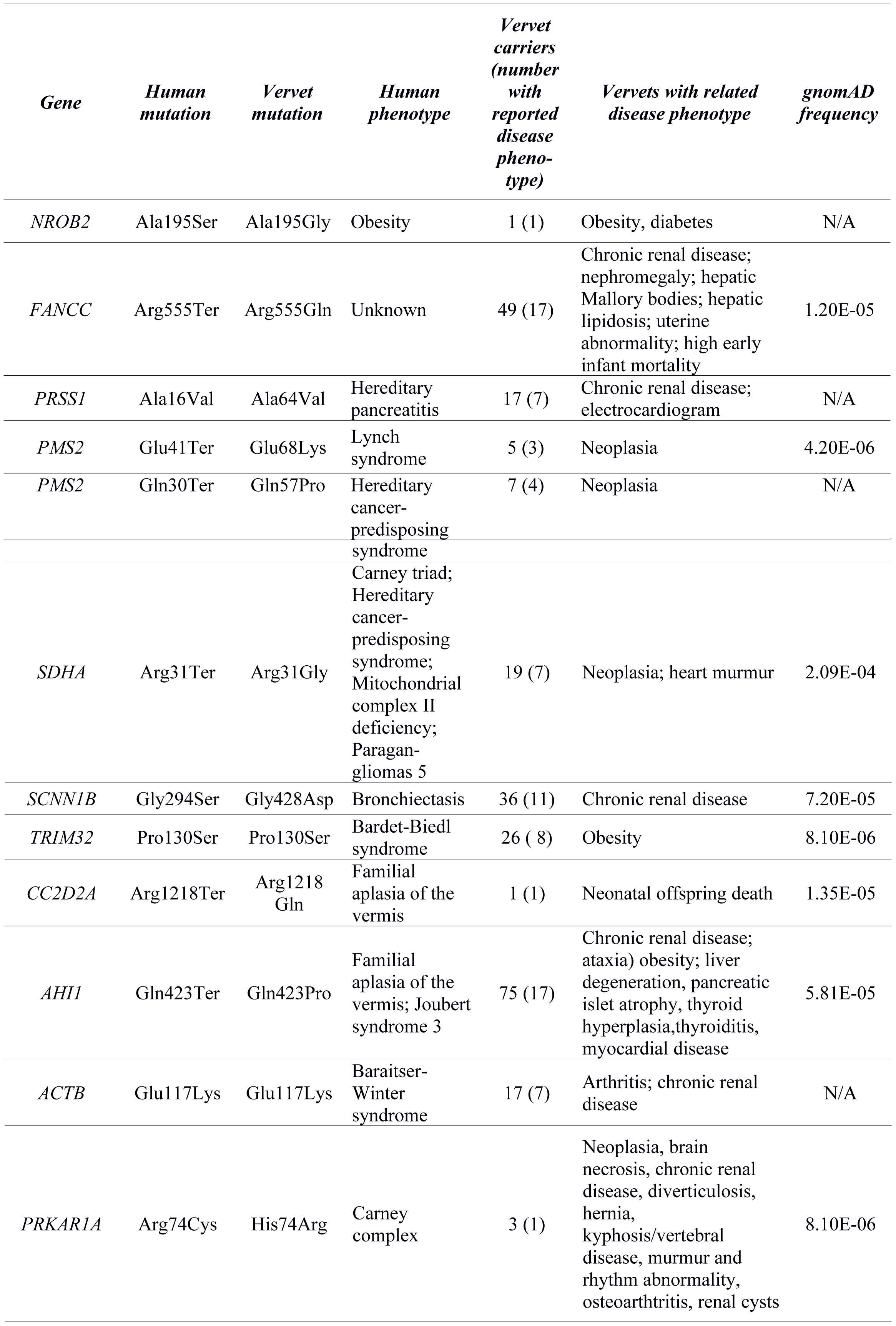
Vervet protein missense variants matching human substitutions with related disease annotation. The protein-coding positional change is noted with the accompanying human disease phenotype. Vervets with recorded phenotypes are provided. If vervet alleles match human we report the allele frequency occurrence based on gnomAD.

## DISCUSSION

In the practice of precision medicine, the accurate filtering of common and rare PTVs of benign consequence remains an unsolved problem. The rapidly expanding repertoire of human sequenced variants classified as putatively damaging has energized discussions on our ability to classify the phenotypic impact of such alleles. Our successful characterization of orthologous vervet genes harboring PTVs offers new opportunities for comparative analyses of PTVs that are either shared between NHPs and humans or private to a species. We report here what is, to our knowledge, the most comprehensive presentation of PTVs transmitted within an NHP biomedical model with ongoing phenotyping. By identifying 2,802 hPTV alleles in 1,747 protein-coding genes that are present in at least one vervet, and reporting the specific monkeys carrying these alleles, we offer the scientific community a starting point for a wide range of further possible studies.

Our initial efforts to relate these PTVs to human phenotypes illustrate these opportunities. We find examples of vervets harboring alternative PTVs that possess clinical features reported in human patients, such as pancreatitis, Bardet-Biedl, and Baraitser-Winter syndromes. For other cases, although the PTVs appear highly penetrant in human, we don’t see comparable phenotypes, motivating research focused on explaining these apparent dichotomies.

Our most extensive evaluation of PTV related vervet phenotypes focused on recurrent early infant mortality. In comparative studies there is often inconclusive evidence for the genetic causes of early infant mortality or miscarriage; for example, homozygous *ARPC3* knockout mice develop only to the blastocyst stage (MGI:1928375), suggesting a higher level of tolerance for such mutations in the vervet. Another promising candidate mutation, occurring in one female vervet for which seven consecutive births resulted in early deaths, was a high-quality splice site variant in hemogloblin, *HBD*. Interestingly, genes that when disrupted are shown to alter blood homeostasis, such as in the thrombophilia condition, are the most popular clinically tested polymorphisms for recurrent miscarriage. Further characterization of the genomes of deceased infants and their mothers will be necessary to close our gaps in understanding the genetic predisposition to miscarriage.

The vervet resource described here is complementary to those generated in other NHP models. Recent sequencing surveys of research colony rhesus macaques have similarly revealed many PTVs that in some cases match human PTVs with relevant clinical data (Xue et al. 2016; Bimber et al. 2017). While in that study, no macaque phenotypes were exploited to correlate to observed phenotypes in human, there is potential to do so in the future. For instance, a rhesus macaque PTV is predicted for *FAM120B*, a gene linked to type I diabetes.

This study provides for the first time, a near-complete survey of variants predicted to disrupt protein-coding within an NHP that has experienced population bottlenecks akin to some human populations. We have catalogued human pathogenic missense and stop-gain mutations from ClinVar (Landrum et al. 2016) that map to polymorphic sites in orthologous vervet genes. We further identified instances in which the alternative allele at such vervet sites is a missense mutation that could potentially contribute to phenotypes with features similar to those observed in human patients. In the future, more extensive searches of human databases that continue to expand the catalog of PTVs, will surely increase the number of vervet PTVs warranting further study.

## METHODS

### Vervet genes

We constructed a comprehensive vervet gene set by merging gene transcripts from NCBI Chlorocebus sabaeus Annotation Release 100 (Pruitt et al. 2012) and ENSEMBL Chlorocebus sabaeus v.1.82 (Aken et al. 2016), excluding pseudogenes. The merged set is non-redundant at the protein level, that is, if two transcripts from NCBI and ENSEMBL translate to the same sequence, only the NCBI transcript is included. Approximately 7% of transcripts have quality issues: missing UTRs (mostly ENSEMBL transcripts), small sequence discrepancies, etc. The set contains 23,250 non-coding transcripts and 67,175 protein-coding transcripts (59,868 NCBI and 7,307 ENSEMBL) that represent 20,533 protein-coding genes. As of Aug. 2016, ENSEMBL and NCBI updated names for ∼1,100 genes, still leaving 3,427 uncharacterized genes not implying orthology with known human genes, for example, LOC103241765 or C1H11orf21.

Vervet protein-coding genes with assigned human orthologs were characterized by recently reported pLI values that reflect probability of a gene of being loss-of-function intolerant (Lek et al. 2016). The analysis of 60,706 high-quality human exome sequences identified 3,230 human genes with pLI values above 0.9 reflecting near-complete depletion of predicted PTV variants (Lek et al. 2016). Such PTV-intolerant genes include virtually all known severe haploinsufficient human disease genes; almost 80% of them have no associated disease phenotype, suggesting undiscovered severe dominant disease genes or genes in which loss of a single copy results in embryonic lethality. Using the known orthologous relationship reflected in the gene names, respective human pLI values were assigned to 15,875 vervet genes out of total 20,080 protein-coding genes with detected sequence variants. Of these, 3,123 genes have pLI>=0.9 (constrained genes) and are thus expected to be more PTV intolerant.

Phenotype associations for human and mouse orthologs of vervet genes were derived from four sources: Mammalian Phenotype Ontology section of MGI resource (Mouse Genome Informatics) (Bult et al. 2016), Human Phenotype Ontology (Köhler et al. 2014), NCBI ClinVar (Landrum et al. 2016) and OMIM (https://omim.org). The meta-list contains gene-phenotype associations for 10,344 human genes; 4,866 genes of this list have at least one annotation related to early infant mortality (EIM): “embryo”, “abnormal reproductive system physiology”, “abnormal survival” sections of the phenotype tree for Mammalian Phenotype Ontology, “abnormality of prenatal development or birth”, “neonatal death”, “stillbirth” for human sources: Human Phenotype Ontology, OMIM and ClinVar.

Protein-protein interactions (PPI) were assessed using the meta-base of STRING (von Mering et al. 2005) and I2D database (Brown and Jurisica 2005) which in turn integrates IntAct, BioGrid, and Mint databases.

### Variant calling and refinement

The BAM files used to call all variant types among the VRC monkeys were as described earlier (Huang et al. 2015). In brief, the average sequence coverage depth varied across the 719 samples in the range 1-39x, with 82 moderate coverage (MC) samples sequenced to the average depth >10x and 637 low coverage (LC) samples with <10x. The sequencing reads in the BAM files were mapped to the pre-submission version of the Vervet Reference Genome, consisting of 2,199 scaffolds. SNV and indel discovery was performed with the Genome Analysis Toolkit (GATK) v3.4 (3.4-0-g7e26428) *HaplotypeCaller* algorithm simultaneously in 719 BAM files over the extended exome regions that included merged gene set exons expanded by +/-50 bp and padded by 1000-bp gene flanks. Total length of the variant calling target is 209 Mb. The called variants with quality QUAL>=20 were lifted over to the *Chlorocebus sabaeus* 1.1 genome assembly using BLAT (Kent 2002). The output contains 1,288,303 biallelic and multiallelic SNVs and short indels. The resulting transition / transversion ratio for biallelic SNVs was 2.45. The sequenced vervet pedigree includes 564 samples in 377 highly overlapping trios that were used to resolve Mendelian errors by refinement of initial genotypes with GATK CalculateGenotypePosteriors (McKenna et al. 2010). Refinement was performed in each trio independently, with results merged to form the consensus genotypes in 719 samples. Low quality genotypes (GQ<10) or genotypes causing Mendelian errors after consensus construction were set to undefined subject in subsequent imputation. This gives 920,346,603 unphased genotypes (1,280,037 sites * 719 samples), approximately 50% of which are undefined due to low coverage depth. WGS processing flowchart is presented in Supplementary Figure 2.

### Genotype phasing and imputation

Genotype phasing and imputation was performed with BEAGLE 4.0 (r1398) software (Browning and Browning 2007, 2009) in two steps. At the first step, the reference set of 82 MC samples and 17 LC samples was phased and imputed. There are 5 trios among the MC samples; the 17 extra samples were selected among the LC samples with <25% of undefined genotypes to provide more trios in the reference set to ensure higher phasing quality. The reference set thus contains 99 samples, among them 5 trios with all MC monkeys and 18 trios with one LC and two MC monkeys. The BEAGLE parameters for burn-in and phasing iterations were set to 5 and 30, respectively.

At the second step, the 620 LC samples were processed in a similar way with 99 samples above used as reference haplotypes. Burn-in, phasing and imputation iteration parameters were set to 5, 20 and 5, respectively. In order to make computations feasible in terms of memory and running time requirements, chromosomes were split into chunks of 3,000 variants overlapping by 500 variants. After completion, consequent overlapping chunks in each sample were “stitched” to form chromosome-wide haplotypes. In the resulting dataset, each genotype was assigned GATK-reported quality of initial genotype (GQ in the range from 0 to 99) and alternative allele dosage DS reported by BEAGLE (DS in the range from 0 to 2). There were three major types of genotypes in the resulting output dataset: (a) called with low quality (GQ<10), set to undefined and imputed; (b) called with intermediate quality (10<=GQ<20) and (c) high quality (GQ>=20). Based on the evaluation of imputation quality (see below), imputed homozygotes genotypes either located at multiallelic sites or assigned dosage parameter in the range 0.1 to 1.9 at biallelic sites were set to undefined because of their low quality. Few genotypes causing Mendelian errors were set to undefined. Sites without homozygous reference alleles were also discarded, leaving in total 896,689,346 genotypes (6.9% of them undefined) at 1,243,738 sites.

### Estimation of imputation quality

To estimate genotype imputation quality, we performed the following test: 10,000 randomly chosen high-quality genotypes (GQ>=50) from chromosomes 6 and 9 in LC samples were set to undefined and imputed along with the other undefined genotypes. Since the overall number of genotypes in the 620 LC samples is approximately 80.5 mln, the additional 10,000 undefined genotypes should not affect the overall imputation quality but instead can be used to measure it. Among the randomly selected genotypes, 3,500 were homozygous and 6,500 heterozygous; 8,280 biallelic and 1,720 multiallelic. After imputation, 3,673 genotypes were called homozygous and 6,327 heterozygous. This observation indicates that, as in the case of called genotypes, the most frequent imputation error type is a heterozygous genotype imputed as homozygous.

Among the 3,673 genotypes imputed as homozygous, the highest error rate was observed for those at multiallelic sites (error rate 38.7%) or biallelic sites with alternative allele dosage DS in the range 0.1 to 1.9, indicating low imputation confidence (error rate 37.2%). Imputed genotypes with such features (homozygotes at multiallelic sites or intermediate dosage at biallelic) comprise 7.6% of the total genotype pool in the 719 samples and were set to undefined based on results of this test. For 3,221 biallelic genotypes imputed as homozygous reference (DS<0.1) or alternative (DS>1.9), error rates are 0.8% and 1.2%, respectively. Out of 6,327 genotypes imputed as heterozygous, 57 (0.9%) are actually homozygous.

Regions of homozygosity were calculated with PLINK v.1.90p (Chang et al. 2015) with the following parameters: `--allow-extra-chr --homozyg extend --homozyg-density 100 --homozyg-kb 100 --homozyg-snp 100 --homozyg-window-het 3 --homozyg-window-snp 50.`

### Variant annotation

Based on location in the gene region and type of nucleotide change, detected SNVs were classified into the following categories: stop-gain, stop-loss, splice-site (donor, acceptor), missense (damaging, benign, unknown), synonymous, UTR exon, non-coding gene exon, gene flank (upstream, downstream), intron. Annotation is non-redundant, that is, if a variant has conflicting annotations in different transcripts of the same gene, the earlier one in the list above was assigned. Classification of missense SNVs into damaging, benign and unknown was performed with PolyPhen-2 (Adzhubei et al. 2010). Indels were classified into the same prioritized categories, excluding stop-gain, stop-loss, missense and synonymous types, replaced by the general “coding exon indel” type.

Our definition of PTVs follows that implemented in the LOFTEE software (Loss-Of-Function Transcript Effect Estimator, https://github.com/konradjk/loftee): stop gain SNVs, splice site disrupting SNVs or indels, frameshifting indels. PTVs do not include indels with length being multiple of 3 (non-frameshifting, or inframe, indels), variants in the last 5% of a transcript, variants in non-canonical and NAGNAG splice sites or surrounding short exons (<15bp). Of 13,777 coding exon indel, splice site or stop gain alleles, 9,574 alleles in 5,668 protein-coding genes passed the PTV criteria above and were used in this study.

### Expression of genes with homozygous PTVs

The available RNA-seq data set reports expression for 33,994 genes in seven tissues of 58 VRC monkeys. The surveyed tissues were adrenal gland, Brodman area 46, blood, caudate nucleus, fibroblasts, hippocampus, and pituitary gland (Jasinska et al. 2017). To evaluate the association of homozygous PTV and harboring gene expression, we used EMMAX software (Kang et al. 2010) with a recessive model for genotype (1 if homozygous for the alternative allele, 0 otherwise). We modeled gene expression as a function of hPTV status with age and sex used as covariates. Expression was inverse-normal transformed prior to analysis. For each PTV we tested expression in each tissue separately. The reported regression coefficient Beta results from modeling expression as a function of genotype and is positive if the alternative homozygous genotype increases gene expression, and negative if it decreases expression.

We considered PTVs that have at least one of 58 animals homozygous for the alternative allele and filtered out genes on sex chomosomes, genes with a mean count of less than 1 across all samples, as well as genes detected in fewer than 10% of samples (see Methods in (Jasinska et al. 2017)). An association test was not performed if expression was constant, or if all animals were of one genotype. Finally, this procedure gave 8,683 tested PTV-tissue combinations for 1,518 PTVs in 1,236 genes (Supplementary Table 4). The stringent set included 3,918 tests for 697 PTVs for which at least three homozygous alternative and three homozygous reference genotypes are high quality genotypes individually (GATK GQ≥20) or belong to runs of homozygosity.

### High early infant mortality and other phenotypes

Among the 719 sequenced VRC monkeys, 272 were females with at least three birth events, for whom there were a total of 1,862 birth events. Of the 272 females with at least 3 births events, we identified 24 females with higher than normal rates of infant mortality (EIM): either infants that were born live and died during the first month of life or were never seen alive due to stillbirths or miscarriage. Using this number, we estimated the overall rate of early infant mortality in VRC as 0.323. Based on the overall EIM rate, we calculated the binomial probability *P* of at least *k* deaths out of *N* total births for each female monkey and identified 18 animals with high EIM rate as *P*<0.05.

Various medical and life trait phenotypes have been reported for single or multiple VRC monkeys, for example, cataract (1 monkey), amyloidosis (3 monkeys), arteriosclerosis (3), arthritis (39), chronic renal disease (27), morphological malformations (3), obesity (62). For the complete list of vervets and phenotypes see Supplementary Table 9.

### Human disease mutations in vervets

We extracted 3,485 sequences of human proteins harboring missense and stop-gain disease mutations annotated as “Pathogenic” in ClinVar and aligned them to all available vervet protein sequences. Alignments with gaps or pairwise identity below 80% were removed, resulting in 22,664 human mutations in 2,810 human proteins. Of these mutations, 67 human mutations mapped to polymorphic vervet sites with missense or stop-gain variants. We kept only variants where reference or both amino acids match, accounting for reported mutation inheritance mode (recessive or dominant). If inheritance mode was not specified, autosomal recessive was assumed. This procedure gave 33 variants that were analyzed using available vervet disease phenotypes and OMIM and Human Phenotype Ontology (Köhler et al. 2014) information. We found 12 mutations where phenotypes observed in vervets harboring alternative allele(s) could be related to clinical features reported in human patients; six instances where human stop-gain variants corresponded to vervet missense variants were discarded, leaving six variants reported (Table 5).

## DATA ACCESS

Underlying sequences used in this study have been deposited in the NCBI Sequence Read Archive (SRA; http://www.ncbi.nlm.nih.gov/sra/) under BioProject accession number PRJNA240242 and in the European Nucleotide Archive under the same accession number (https://www.ebi.ac.uk/ena/data/view/PRJNA240242) (Huang et al. 2015). Users are advised to enable the “Submitter’s sample name” column in the selection table to ensure display of vervet IDs used in this paper. Individual genotypes are available via vervet genome browser at https://coppolalab.ucla.edu/vgb/home.

## ACKNOWLEDGEMENTS

Funding to Richard K Wilson was provided by NIH-NHGRI grant 5U54HG00307907. Funding to Jay R Kaplan was provided by P40RR019963/OD010965. Funding to Nelson B. Freimer was provided by NIH grants R01RR016300 and R01OD010980. Funding to Vasily Ramensky was provided by RFBR, research project No. 18-04-00789 A.

## AUTHOR CONTRIBUTIONS

WCW, MJ, JK, MC, and RKW, produced the data. VR, AJJ, AZ, NBF, SKS, WCW analyzed the data. VR, NBF, AJJ, and WCW designed the study. VR, WCW and NBF wrote the paper. All authors reviewed and approved the final draft.

