## Supplemental Fig. 1 for "The burden of deleterious variants in a non-human primate biomedical model"

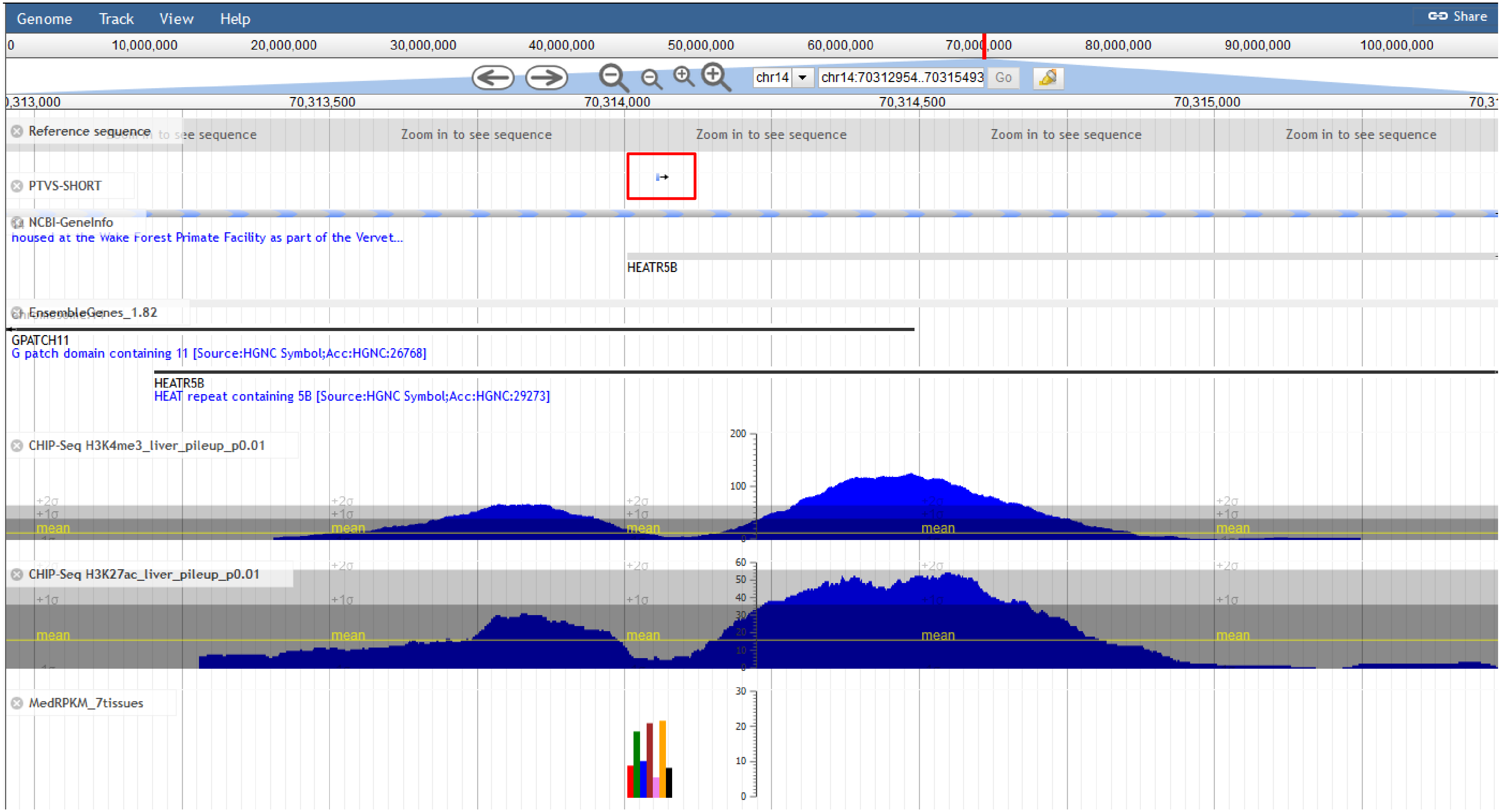
Genome
Track
View
Help
Share
chr14
chr14:70312954..70315493
Go
Reference sequence
Zoom in to see sequence
PTVS-SHORT
NCBI-GeneInfo
housed at the Wake Forest Primate Facility as part of the Vervet...
HEATR5B
EnsembleGenes_1.82
GPATCH11
G patch domain containing 11 [Source:HGNC Symbol;Acc:HGNC:26768]
HEATR5B
HEAT repeat containing 5B [Source:HGNC Symbol;Acc:HGNC:29273]
CHIP-Seq H3K4me3_liver_pileup_p0.01
CHIP-Seq H3K27ac_liver_pileup_p0.01
MedRPKM_7tissues

(A)

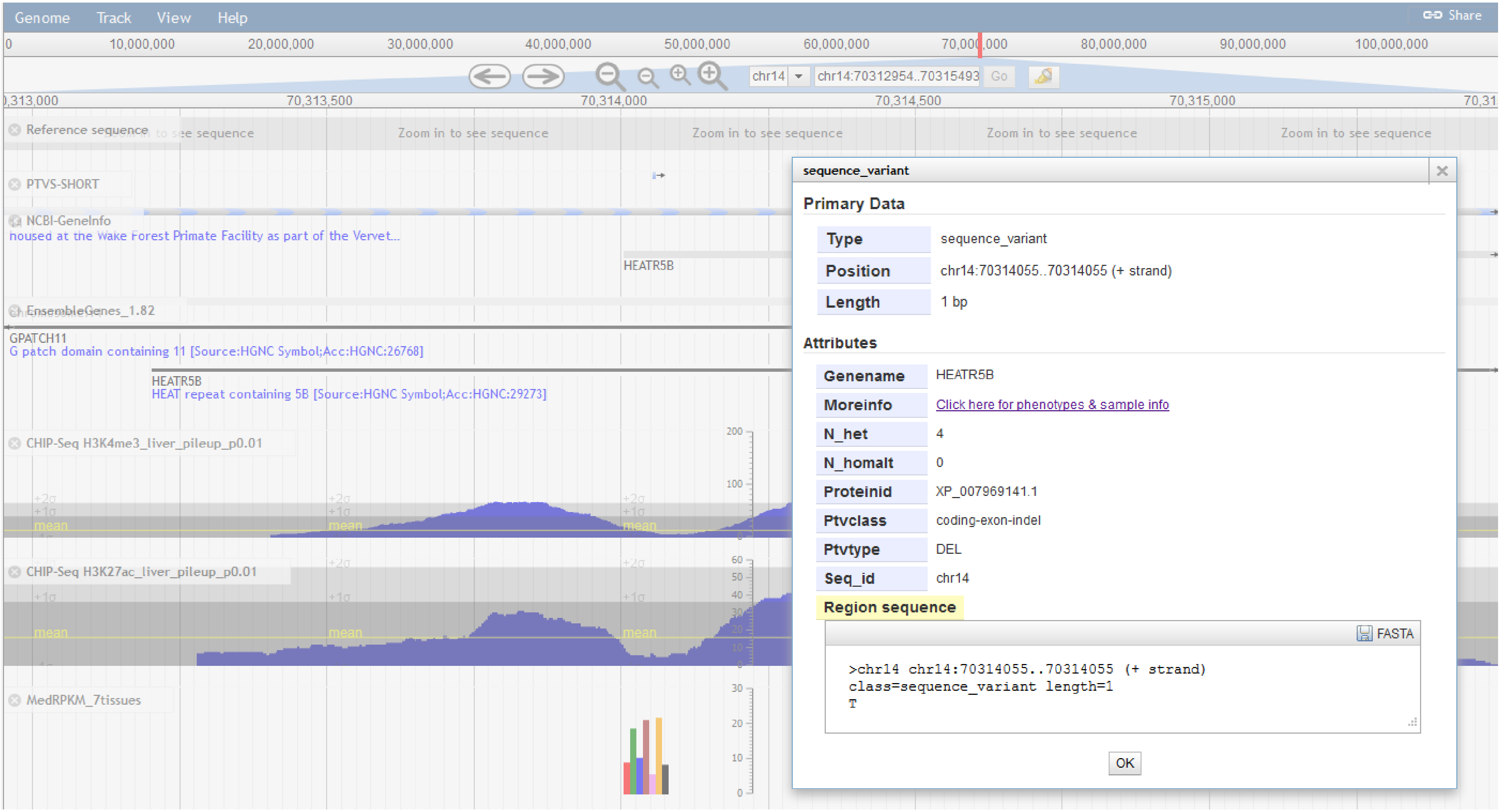
Genome
Track
View
Help
Share
chr14
chr14:70312954..70315493
Go
Reference sequence
Zoom in to see sequence
PTVS-SHORT
NCBI-GeneInfo
housed at the Wake Forest Primate Facility as part of the Vervet...
HEATR5B
EnsembleGenes_1.82
GPATCH11
G patch domain containing 11 [Source:HGNC Symbol;Acc:HGNC:26768]
HEATR5B
HEAT repeat containing 5B [Source:HGNC Symbol;Acc:HGNC:29273]
CHIP-Seq H3K4me3_liver_pileup_p0.01
CHIP-Seq H3K27ac_liver_pileup_p0.01
MedRPKM_7tissues
sequence_variant
Primary Data
Type
sequence_variant
Position
chr14:70314055..70314055 (+ strand)
Length
1 bp
Attributes
Genename
HEATR5B
Moreinfo
Click here for phenotypes & sample info
N_het
4
N_homalt
0
Proteinid
XP_007969141.1
Ptvclass
coding-exon-indel
Ptvtype
DEL
Seq_id
chr14
Region sequence
FASTA
>chr14 chr14:70314055..70314055 (+ strand)
class=sequence_variant length=1
T
OK

**(B)**

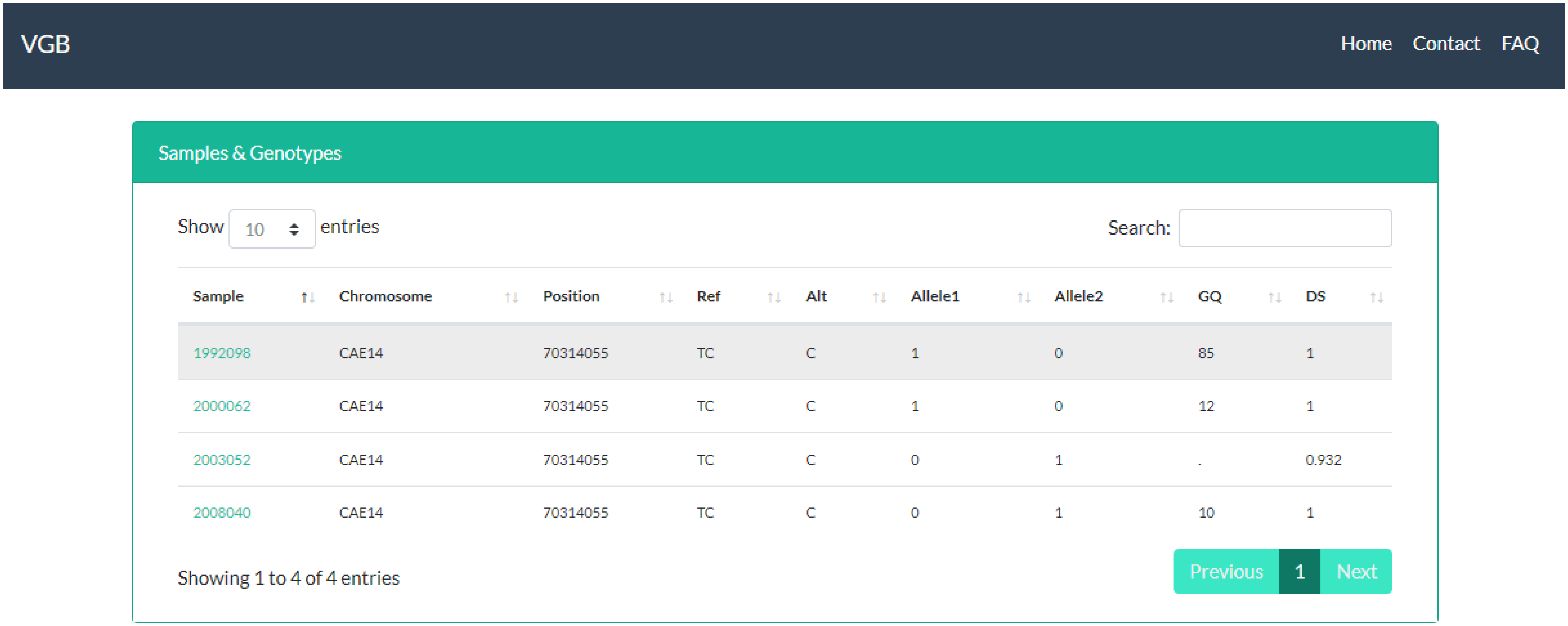


| Sample | Chromosome | Position | Ref | Alt | Allele1 | Allele2 | GQ | DS |
|---|---|---|---|---|---|---|---|---|
| 1992098 | CAE14 | 70314055 | TC | C | 1 | 0 | 85 | 1 |
| 2000062 | CAE14 | 70314055 | TC | C | 1 | 0 | 12 | 1 |
| 2003052 | CAE14 | 70314055 | TC | C | 0 | 1 | . | 0.932 |
| 2008040 | CAE14 | 70314055 | TC | C | 0 | 1 | 10 | 1 |

**(C)**

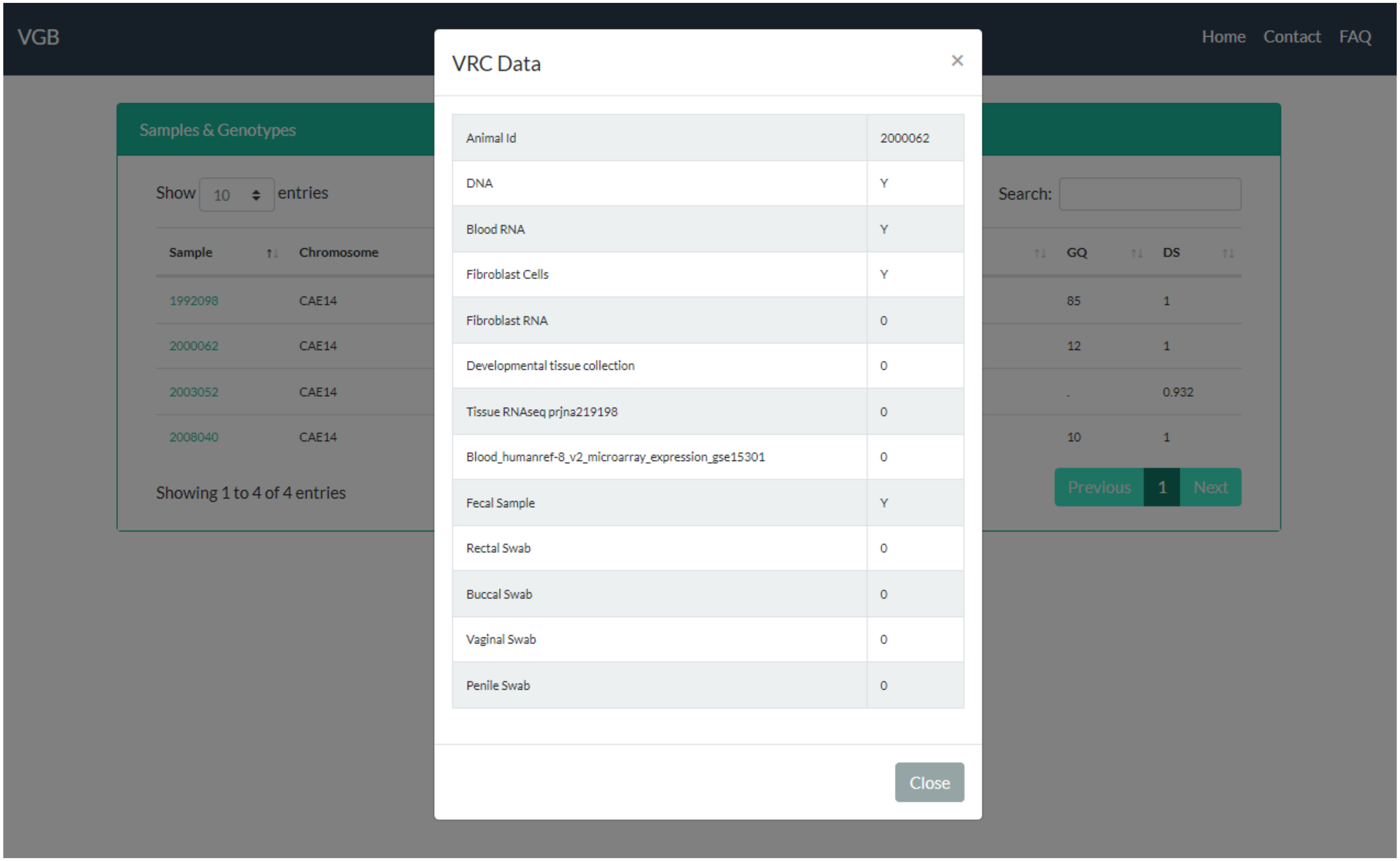


**(D)**

Supplementary Figure 1. Verver genome browser. (A) Major tracks with PTV of interest denoted by red rectangle. (B) PTV information pop-up window. (C) Genotype information for PTV carriers. (D) Biological sample availability for a PTV carrier.
