## Supplemental Fig. 2 for "The burden of deleterious variants in a non-human primate biomedical model"

Non-redundant set of NCBI and ENSEMBL genes:
*23,250 non-coding and 67,175 coding transcripts*

*719 BAM files (Huang 2015): 82 moderate coverage (MC) samples and 637 low coverage (LC)*

↓

Joint SNP and indel calling in extended exome regions in all samples (GATK HaplotypeCaller)

↓

Lift over to the Chlorocebus sabaeus 1.1 genome assembly.
*1,288,303 SNVs and indels with QUAL>=20*

↓

Mendelian errors in trios resolved (GATK CalculateGenotypePosteriors).
Genotypes with GQ<10 and remaining errors set to undefined.
*920,346,603 unphased genotypes.*

↓

82 medium coverage (MC) and 17 low coverage (LC) samples phased and imputed (BEAGLE 4.0)

↓

620 LC samples phased and imputed with 99 MC and LC used as reference. Haplotype fragments stitched. Estimation of imputation quality.

↓

Low quality imputed homozygotes genotypes set to undefined. Regions of homozygosity were calculated (PLINK v.1.90p).
*896,689,346 genotypes (6.9% undefined) at 1,243,738 sites.*

↓

Variant annotation (LOFTEE, PolyPhen-2).
*9,574 protein-truncating variants in 5,668 protein-coding genes.*

Supplementary Figure 2. Flowchart of variant calling, imputation and annotation.
